# Structural basis of GIRK2 channel modulation by cholesterol and PIP_2_

**DOI:** 10.1101/2020.06.04.134544

**Authors:** Yamuna Kalyani Mathiharan, Ian W. Glaaser, Yulin Zhao, Michael J. Robertson, Georgios Skiniotis, Paul A. Slesinger

## Abstract

G protein-gated inwardly rectifying potassium (GIRK) channels play important roles in controlling cellular excitability in the heart and brain. While structural data begin to unravel the molecular basis for G protein and alcohol dependent activation of GIRK channels, little is known about the modulation by cholesterol. Here, we present cryo-electron microscopy (cryoEM) structures of GIRK2 in the presence and absence of the cholesterol analog cholesteryl hemisuccinate (CHS), and PIP_2_. The structures and their comparison reveal that CHS binds near PIP_2_ in lipid-facing hydrophobic pockets of the transmembrane domain (TMD). CHS potentiates the effects of PIP_2_, which stabilizes the inter-domain region and promotes the engagement of the cytoplasmic domain (CTD) onto the transmembrane region. The results suggest that CHS acts as a positive allosteric modulator and identify novel therapeutic sites for modulating GIRK channels in the brain.

---

G protein-gated inwardly rectifying potassium (GIRK) channels provide a major source of inhibition in the brain, and have been implicated in a variety of neurological disorders^1^. Loss of GIRK channels leads to hyperexcitability and susceptibility to seizures, changes in alcohol consumption, and increases in sensitivity to psychostimulants^2–6^. Activation of G protein-coupled receptors (GPCRs) that signal through Gα_i/o_ G protein, such as GABA_B_, D2 dopamine and m2 muscarinic receptors, leads to liberation of G protein Gβγ subunits that open GIRK channels through direct protein-protein interactions^7–9^. GIRK channels are also activated directly by alcohol^10–12^. Recently, GIRK channels have been shown to be modulated by membrane cholesterol^13–15^, similar to other ion channels such as calcium-activated potassium channels (‘BK”) and TRP channels^15,16^. GIRK currents are significantly larger in *Xenopus* oocytes enriched with cholesterol^14,15^ and, in a minimal lipid vesicle reconstitution experiment with only GIRK2 and PIP_2_, the addition of cholesterol significantly enhances potassium conductance^13^.

Some of the questions on the molecular aspects of GIRK channel gating have been addressed through a combination of electrophysiology, biochemistry, mutagenesis and determination of high-resolution structures. For example, Gβγ binds directly to the channel via the βL-βM loop in the cytoplasmic domain (CTD)^17–20^. The binding induces a conformational change in the channel’s two gates, the G loop gate^21^ and the helix bundle crossing (HBC) gate^22^, widening the pore to enable K^+^ permeation, as suggested by the X-ray crystal structures of GIRK2 in complex with Gβγ (PDBID:4KFM)^18^ and GIRK2 containing a mutation (R201A) meant to mimic the G protein activated state (PDBID:3SYQ)^23^. A physical binding pocket for mediating alcohol-dependent activation has been also localized in the CTD, formed by the three domains at the subunit interface, the βL-βM loop from one subunit and the βD-βE loop and part of the N-terminal domain from the other subunit^11,24^. The activation of GIRK channels by Gβγ and alcohol both require the membrane phospholipid PIP_2_, which interacts with a cluster of basic residues located at the inner membrane leaflet near the cytoplasmic side of the M1 and M2 transmembrane helices^23,25^. The mechanism by which PIP_2_ supports channel activation, however, is poorly understood, partly due to structural differences observed in the apo state (i.e., no PIP_2_) of inward rectifier channels^23,26^.

Moreover, the mechanism of modulation of GIRK channels by cholesterol is unclear as no binding sites have been directly determined by structural studies. Cholesterol was originally hypothesized to alter the function of proteins in the plasma membrane through changes in membrane stiffness^27,28^. However, more recent studies suggest that cholesterol exerts specific effects through direct protein engagement^27,28^. Putative cholesterol-binding motifs (e.g., CRAC, CARC, CCM) have been identified in some proteins (for review, see^27^). Consistent with this, several structures of GPCRs have shown cholesterol bound to a hydrophobic region of the receptor^27,29–33^. Mutagenesis studies and simulations have implicated several different regions of inwardly rectifying potassium channels in cholesterol modulation, including the cytoplasmic domain, PIP_2_ binding site, and the transmembrane domain^14,34–37^. However, the exact location of the cholesterol sites remains to be determined.

To better understand the structural basis for cholesterol and PIP_2_ modulation of GIRK channels, we employed cryo-electron microscopy (cryoEM) to visualize GIRK2 under different conditions. Here, we present structures of the GIRK2 channel in the presence and absence of the cholesterol derivative cholesteryl hemisuccinate (CHS) and PIP_2_, revealing their effects on mechanistic aspects of GIRK function.

## RESULTS

### CHS potentiates GIRK2 channel activity

Previously, we showed that GIRK2 channels are potentiated by cholesterol in the absence of G proteins^13^. To better understand the mechanism of cholesterol modulation, we sought to determine the site of interaction of a cholesterol analog, cholesteryl hemisuccinate (CHS), in GIRK2 channels. To verify the functional effects of CHS, we employed *Mus musculus* GIRK2 recombinantly expressed in *Pichia pastoris* and extracted from membranes with n-Dodecyl-D-β maltoside (DDM) detergent in the presence or absence of CHS (**Extended Fig. S1a**). Purified GIRK2 tetrameric channels were reconstituted into liposomes containing 1% brain PIP_2_ alone, which is required for channel function^13,38^, and either 2.5% CHS or 5% cholesterol. We measured K^+^ permeation through GIRK2 channels using a flux assay (**Fig. 1a**), as described previously^13,18^. Upon addition of CCCP, the fluorescence emitted from ACMA pre-loaded into proteoliposomes containing GIRK2 channels and 1% brain PIP_2_ (GIRK2^PIP2^) quenches to ~ 0.5 F/F_0_ under basal conditions (**Fig.1a and b**). Next, we reconstituted membranes in CHS (GIRK2^PIP2/CHS^) and observed fluorescence quenching increasing to ~ 0.2 F/F_0_ following CCCP, indicating potentiation of GIRK2 channel activity **(Fig.1a and b).** The GIRK2 channel-specific inhibitor, MTS-HE^13^, reduced the extent of quenching, indicating inhibition of the K^+^ conductance. We used the inhibition with MTS-HE to quantify changes in quenching under different conditions and converted this to a percentage of relative K^+^ flux. Under basal conditions with proteoliposomes containing GIRK2 channels and 1% brain PIP_2_ (GIRK2^PIP2^), the % relative K^+^ flux was 44.0% ± 2.3% (n=18) (**Fig. 1c-e**). Both CHS and cholesterol significantly increased the relative K^+^ flux to 65.2% ± 0.4% (n=10) and 69.4% ± 0.3% (n=17), respectively, compared to basal conditions (**Figure 1c-e**). Taken together, these results demonstrate that CHS, like cholesterol, potentiates basal PIP_2_-dependent GIRK2 channel activity^13^.

**Figure 1:**
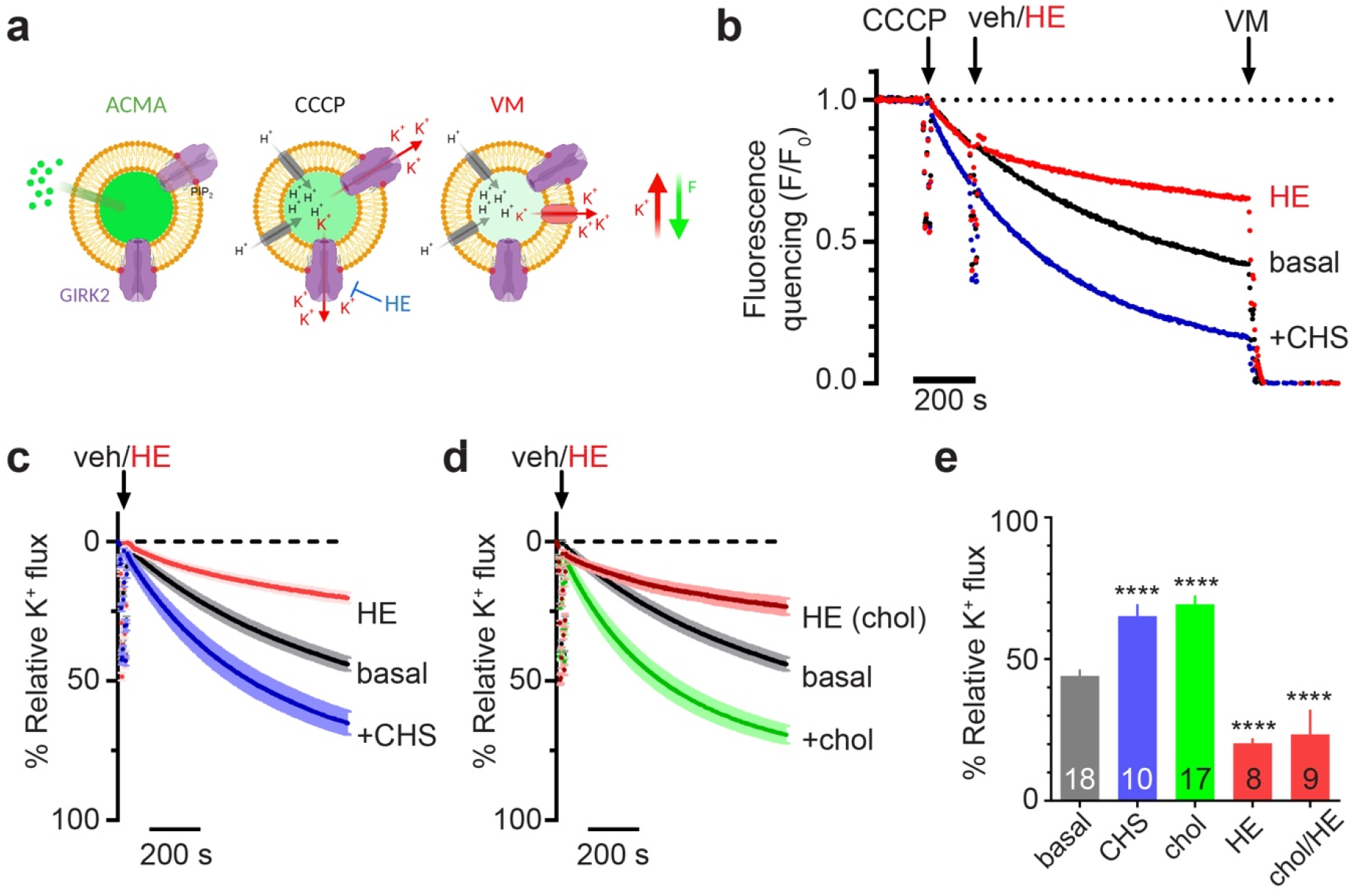
CHS potentiates GIRK2 activity in proteoliposomes with PIP_2_. **(a)** Schematic of the K^+^ flux assay. Left panel indicates proteoliposomes pre-loaded with ACMA at baseline. Middle panel demonstrates fluorescence quenching in the presence of open GIRK channels upon addition of CCCP and H^+^ influx. Right panel shows the dissipation of the K^+^ gradient upon addition of valinomycin (VM), indicating the vesicle capacity. **(b)** Example of full-length normalized fluorescent traces of GIRK2-containing liposomes with 1% brain PIP_2_. Fluorescence quenching is measured upon addition of CCCP (arrow), and then after addition of GIRK2 channel inhibitor 100 μM MTS-HE (HE) or vehicle (veh, arrow), and then terminated by addition of VM. Note increase in quenching in the presence of CHS (blue trace), as compared to basal (black trace) and HE inhibited (red trace). **(c)** Plot shows normalized % relative K^+^ flux (mean ± SEM) of GIRK2-containing liposomes with 1% brain PIP_2_ + CHS (blue trace, n=10) or 1% brain PIP_2_ alone (basal; black trace, n=18) following addition of veh, or after the addition of MTS-HE for 1% brain PIP_2_ alone (red trace, n=8). **(d)** Normalized % relative K^+^ flux (mean ± SEM) of GIRK2-containing liposomes with 1% brain PIP_2_ + 5% cholesterol (green trace, n=17) or 1% brain PIP_2_ alone (black, same as b) following addition of veh, or following addition of HE for 1% brain PIP_2_ + 5% cholesterol (red trace, n=9). **e)** Bar graph shows the average (±◻SEM; n indicated on graph) % relative K^+^ flux for the indicated conditions. Both CHS and cholesterol significantly increased the % relative K^+^ flux (**** p<0.0001 vs. basal). HE (red bar) significantly inhibited GIRK2 channels.

### Structure of GIRK2 in the presence of CHS and PIP_2_

To gain insights to CHS binding in GIRK2, we proceeded with optimizing the sample for cryoEM studies. The sample quality was assessed by size-exclusion chromatography (**Extended Fig. S1a**) and further evaluated by negative stain electron-microscopy (**Extended Fig. S1b)** to identify conditions with predominantly well-formed GIRK2 tetramers^39^. To ensure the presence of CHS and PIP_2_ (GIRK2^PIP2/CHS^), the sample was purified in the presence of CHS and incubated with 2 mM of diC_8_-PIP_2_ prior to cryoEM grid preparation.

We subsequently employed cryoEM and determined the structure of GIRK2^PIP2/CHS^ at a global indicated resolution of 3.5 Å (**Extended Fig. S2**). The structure revealed an overall architecture similar to those solved by X-ray crystallography^18,23^, with two distinct domains, the transmembrane (TMD) and cytoplasmic (CTD), and the pore along the four-fold axis with well-ordered inner-helix bundle crossing and G-loop gates (**Fig. 2a and 2b**). PIP_2_ is bound at its canonical site near inter-domain linkers, as also observed in other inwardly rectifying potassium channels^18,23,26^ (**Fig. 2b).** Interestingly, we observed a planar density on the opposite side of the PIP_2_-channel interface that is consistent with CHS, hereafter referred to as CHS1. Initial modeling of CHS in the EM density was performed with the GemSpot pipeline^40^, followed by manual refinement in COOT^41^. This approach yielded largely identical poses for CHS in good agreement to the EM map and chemically plausible interactions with GIRK2 transmembrane helices that had subtle differences in the orientation of iso-octyl tail (**Fig. 2b-d and Extended Fig. S3a**). The head group of CHS1 is in position to form a salt bridge with R92, while its sterane rings and iso-octyl tail are stacked against the TMD, surrounded by hydrophobic residues F93, L95, L96, V99 of M1 helix from one subunit, and I175, V178, L179 of M2 helix from the adjacent subunit (**Fig. 2b-d and Extended Fig. S3b**). Another CHS molecule (CHS2) could be modelled near the N-terminus surrounded by hydrophobic residues that include V72, L79, I82, L86, L89, I97, V101 and F186 (**Fig. 2b-d and Extended Fig. S3c**). We cannot rule out other putative CHS sites, as additional densities are observed near the TMD-CTD interface, but those are poorly resolved (**Extended Fig. S3d**).

**Figure 2:**
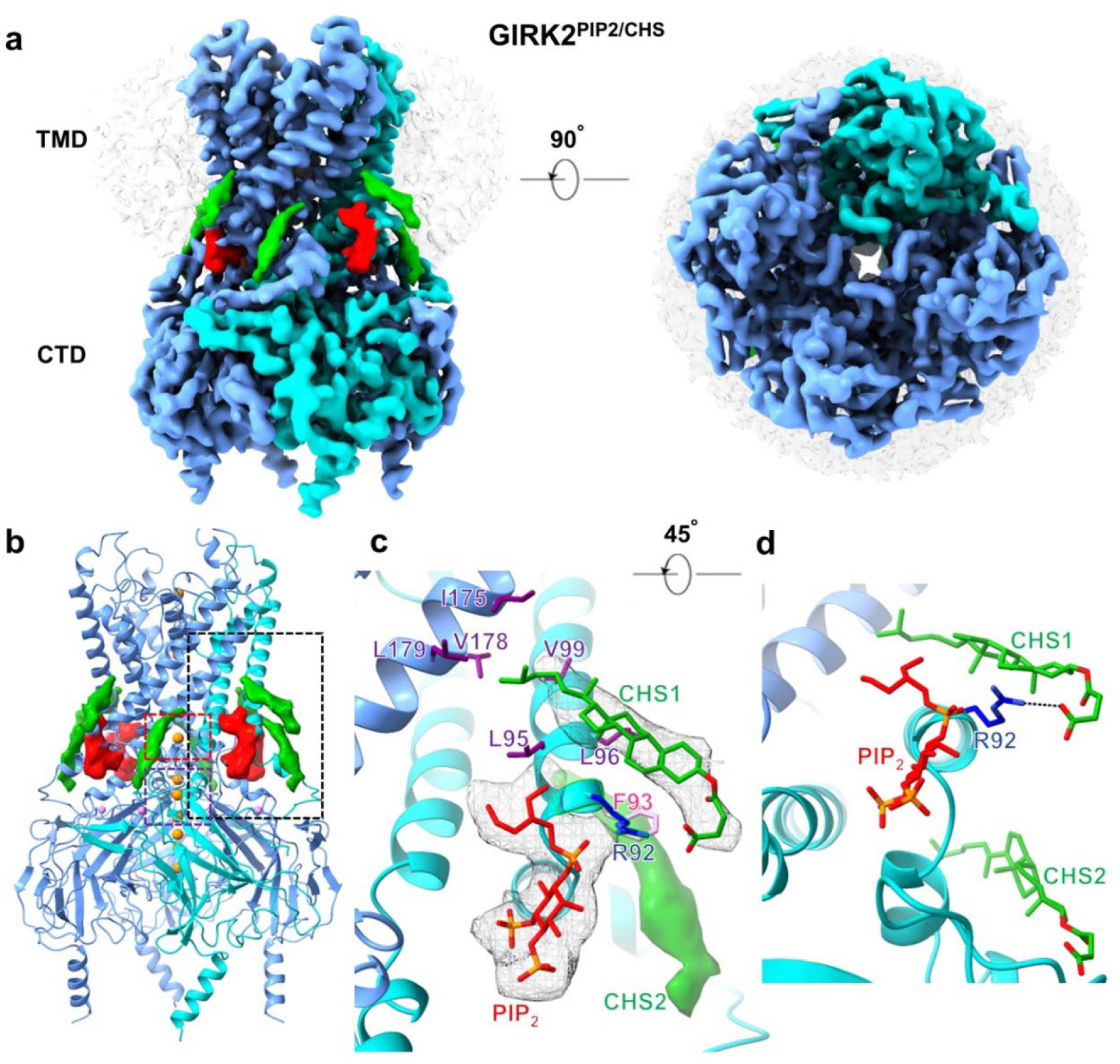
Structure of GIRK2 with PIP_2_ and cholesteryl hemisuccinate (CHS). **(a)** GIRK2^PIP2/CHS^ cryoEM map; two CHS (green) molecules per subunit are modelled into planar densities on the opposite side of PIP2 (red)-channel interface. GIRK2 subunits are in blue with one highlighted in cyan; DDM micelle is shown in transparency. The top right panel shows bottom-top view from cytoplasmic side. **(b)** Model of GIRK2^PIP2/CHS^ with the transmembrane and cytoplasmic domains. K^+^ ions (orange) are shown along the pore and one Na^+^ ion (violet) is modelled per subunit as seen in X-ray crystal structures. The densities of CHS and anionic head group of PIP_2_ are shown at the TMD-CTD interface. The inner-helix and G-loop gates are highlighted in red and purple dashed boxes, respectively, and observed CHS pockets is in black dashed box. **(c)** CHS (green, density shown as mesh, referred to as CHS1) modeled into a density near PIP_2_ (red, density shown as mesh) is surrounded by positively charged R92 (dark blue) and hydrophobic residues including F93 (pink), L95, L96, V99 (purple) from one protomer (cyan) and I175, V178, L179 (purple) from nearby subunit (blue). The density of other CHS molecule (green, referred to as CHS2) adjacent to this is represented as surface. The densities of anionic head group of PIP_2_ and CHS are contoured at a threshold level of 0.0073 in ChimeraX. **(d)** CHS1 is on the opposite side of PIP_2_ in the GIRK2^PIP2/CHS^ map and its carboxylate group coordinates with the side chain of R92 (dark blue).

### Structures of GIRK2^PIP2^ in the absence of CHS

To assess whether CHS alters the interaction of PIP_2_, we next determined the structure of GIRK2^PIP2^ in the absence of CHS during the purification. Interestingly, cryoEM analysis resolved different conformers of GIRK2^PIP2^ (**Extended Fig. S4**). The predominant conformer, referred to as GIRK2^PIP2^*, accounted for ~ 39% of well-defined particles and its structure was obtained at a global indicated resolution of 3.2 Å. This structure is comparable to GIRK2^PIP2/CHS^ (**Fig. 3**), with four PIP_2_ molecules bound at equivalent sites, but lacked densities for CHS1 and CHS2 (**Extended Fig. 3d**). A second conformer, GIRK2^PIP2^**, accounting for ~ 27% of projections was determined at 4.8 Å global resolution. Comparison of this conformation with GIRK2^PIP2^* reveal that the CTD is twisted from 8-12° around the four-fold axis and is partially disengaged from the TMD by 1-2 Å (as evaluated by the distance between C α atoms of I195 and Q322), with a stretched tether helix (residues 197-203, **Extended Fig. S5a-c**).

**Figure 3:**
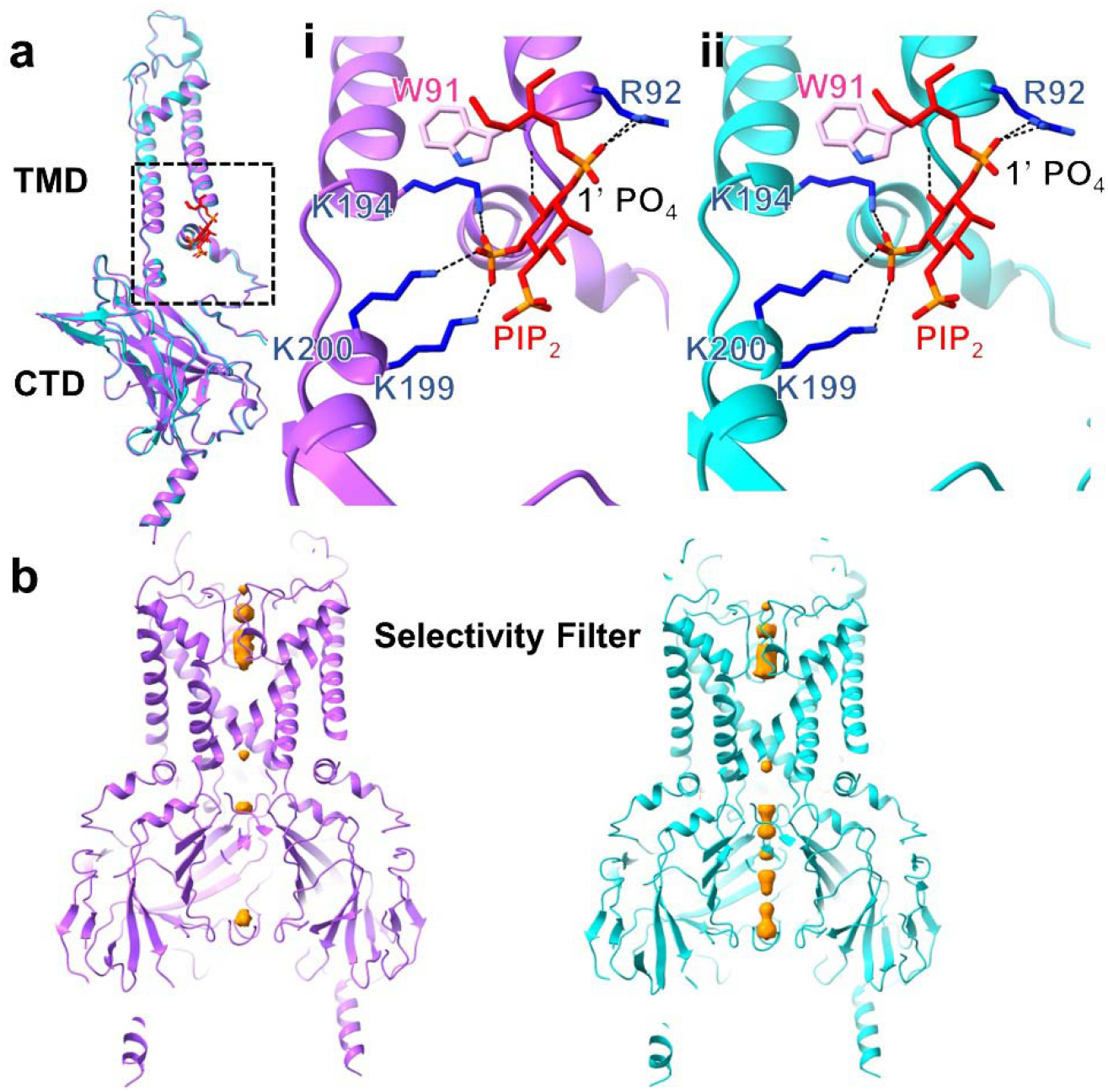
PIP_2_ binding pocket and K^+^ ions present in cryoEM structures. **(a)** Comparison of a subunit from GIRK2^PIP2^* (purple) and GIRK2^PIP2/CHS^ (cyan) structures; the RMSD between them is around 0.4 Å. Insets **(i)** and **(ii)** represent PIP_2_ (red) coordination in GIRK2^PIP2^* (purple) and GIRK2^PIP2/CHS^ (cyan) respectively; PIP_2_ interacts with side chain of positively charged residues R92, K194, K199, K200 (dark blue) and amide backbone of W91 (pink). In cryoEM structures, the 1’ PO_4_ of PIP_2_ is coordinated by the side chain of R92. **(b)** K^+^ ions were modelled into densities in the pore of GIRK2^PIP2^* (purple) and GIRK2^PIP2/CHS^ (cyan) cryoEM maps (orange, contoured at a threshold level of 0.0045 in ChimeraX). Apart from the selectivity filter, additional densities compatible with K^+^ ions are observed along the pore.

The disengagement of the CTD is more pronounced in two additional conformers, GIRK2^PIP2^*** and GIRK2^PIP2^****, as the distance between Cα atoms of I195 and Q322 is increased by 3-8 Å (**Extended Fig. S5a and b**). These conformers appear very similar and were obtained at low overall resolution (~7.7 Å), primarily due to the relative flexibility between TMDs and CTDs, but nevertheless rigid body docking was sufficient in revealing the overall backbone configurations. Interestingly, though the CTD detachment is different in each subunit, its C4 symmetry appears nearly preserved in all the conformers. We next examined whether the interaction of PIP_2_ in its binding pocket was altered by the presence of CHS (**Fig. 3a**). The 1’ PO_4_ of the PIP_2_ anionic headgroup is coordinated by the side chain of positively charged R92 while the 5’ PO_4_ with K194, K199, K200, and O6 atom with the backbone amide group of W91 (**Fig. 3ai**). Positively charged residues K64 and K90 are also in proximal distance to the anionic headgroup of PIP_2_. This configuration of PIP_2_ looks similar to that in GIRK2^PIP2/CHS^ (**Fig. 3aii**), but has lower occupancy. Taken together, these observations suggest that the inter-domain interface of GIRK2^PIP2^ is relatively less stable than that of GIRK2^PIP2/CHS^ (**Extended Fig. S5a and b**).

In both the GIRK2^PIP2^* and GIRK2^PIP2/CHS^ maps, we observe densities that are compatible with Na^+^ and K^+^ ions. A Na^+^ ion is coordinated by D228, as shown previously in X-ray crystal structures^23^. Similarly, densities for K^+^ ions are found near the selectivity filter, Y266, M319, G318 and E236 as previously determined^18,23^ but also observed at new positions near G158, F192 and M313 (**Fig. 3b**). We also note the presence of additional, less well resolved densities along the pore near N184, V188, E236 and M319. Thus, the channel with four PIP_2_ molecules bound at the TMD-CTD appears to be in a state favorable for conducting K^+^ ions through the pore.

### Apo GIRK2 structure (no PIP_2_)

Lastly, we examined the structure of apo GIRK2, in the absence of modulators PIP_2_ and CHS. CryoEM maps of apo GIRK2 determined at overall 4.8 Å resolution revealed that the CTD is detached from the TMD (**Fig. 4 and 5**). This configuration of the CTD is quite different from the apo GIRK2 structure determined by X-ray crystallography, which showed the CTD engaged with the TMD^23^ (**Extended Fig. S5d**). Inter-domain linkers consisting of tether helix and N-terminal residues 67-78 appear mostly disordered, suggesting that this interface is unstable in the absence of PIP_2_ and CHS (**Fig. 5; Extended Fig. S5e, S6 and Extended Movie S1**). The gap between the CTD and TMD is 23-28 Å (**Fig. 4**). The detached CTD appears tilted below the membrane, reflecting ‘wobble’ in this structure and an overall asymmetry in the absence of PIP_2_ and CHS. Thus, our cryoEM studies reveals a new apo GIRK2 closed state.

**Figure 4:**
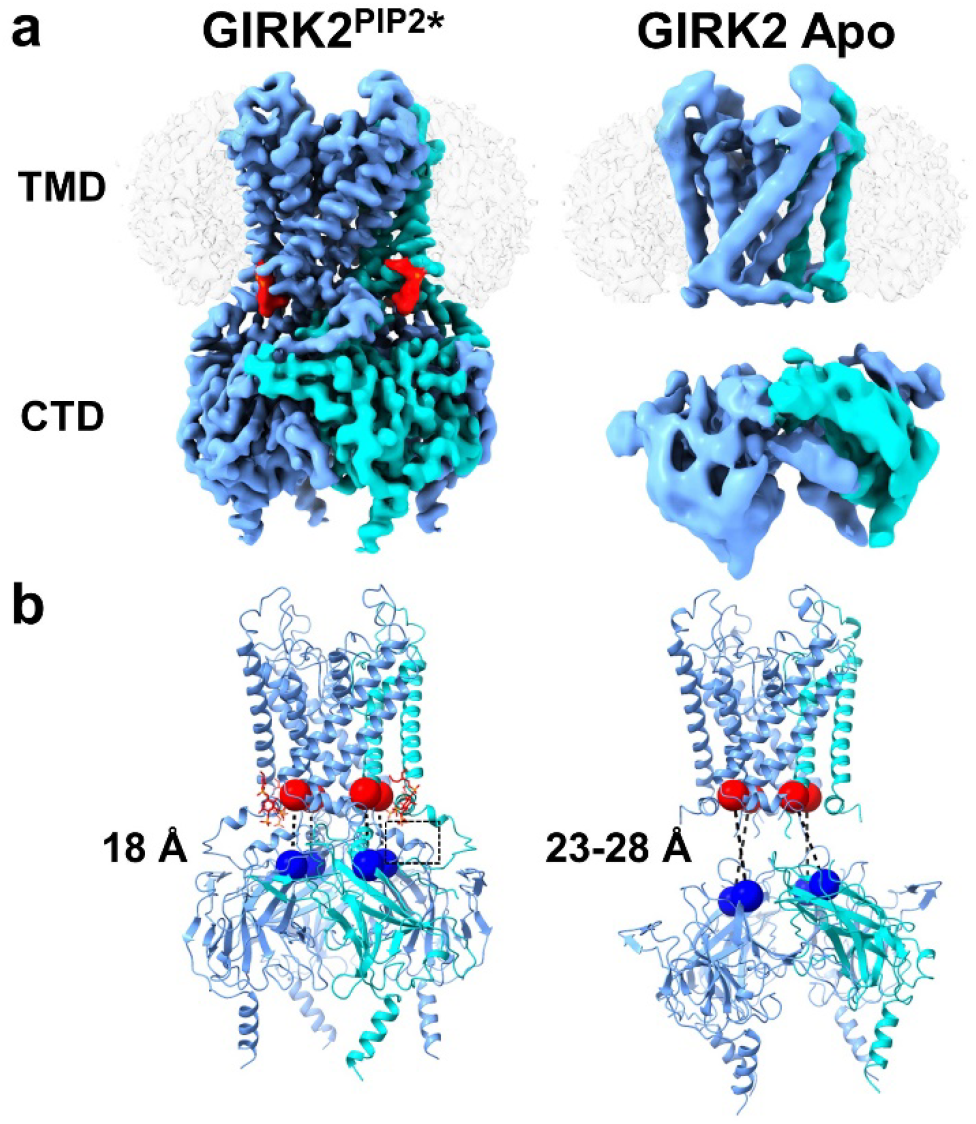
CryoEM structures of GIRK2 with and without PIP_2_. **(a)** CryoEM maps of the most prevalent GIRK2^PIP2^* and GIRK2 apo conformations; the CTD is disengaged from membrane in apo compared to the GIRK2^PIP2^* structure with the tether helix (residues 197-203) and N-terminal residues 67-78 disordered. PIP_2_ (red) is at TMD-CTD interface, one subunit is highlighted in cyan and micelle is in transparency. **(b)** Evaluation of CTD detachment in GIRK2^PIP2^* and GIRK2 apo, the distance between Cα atoms of residues I195 (denoted by red sphere) and Q322 (denoted by blue sphere) in these structures are 18 and 23-28 Å, respectively. The ordered tether helix from one subunit of GIRK2^PIP2^* is shown with a dashed square.

**Figure 5:**
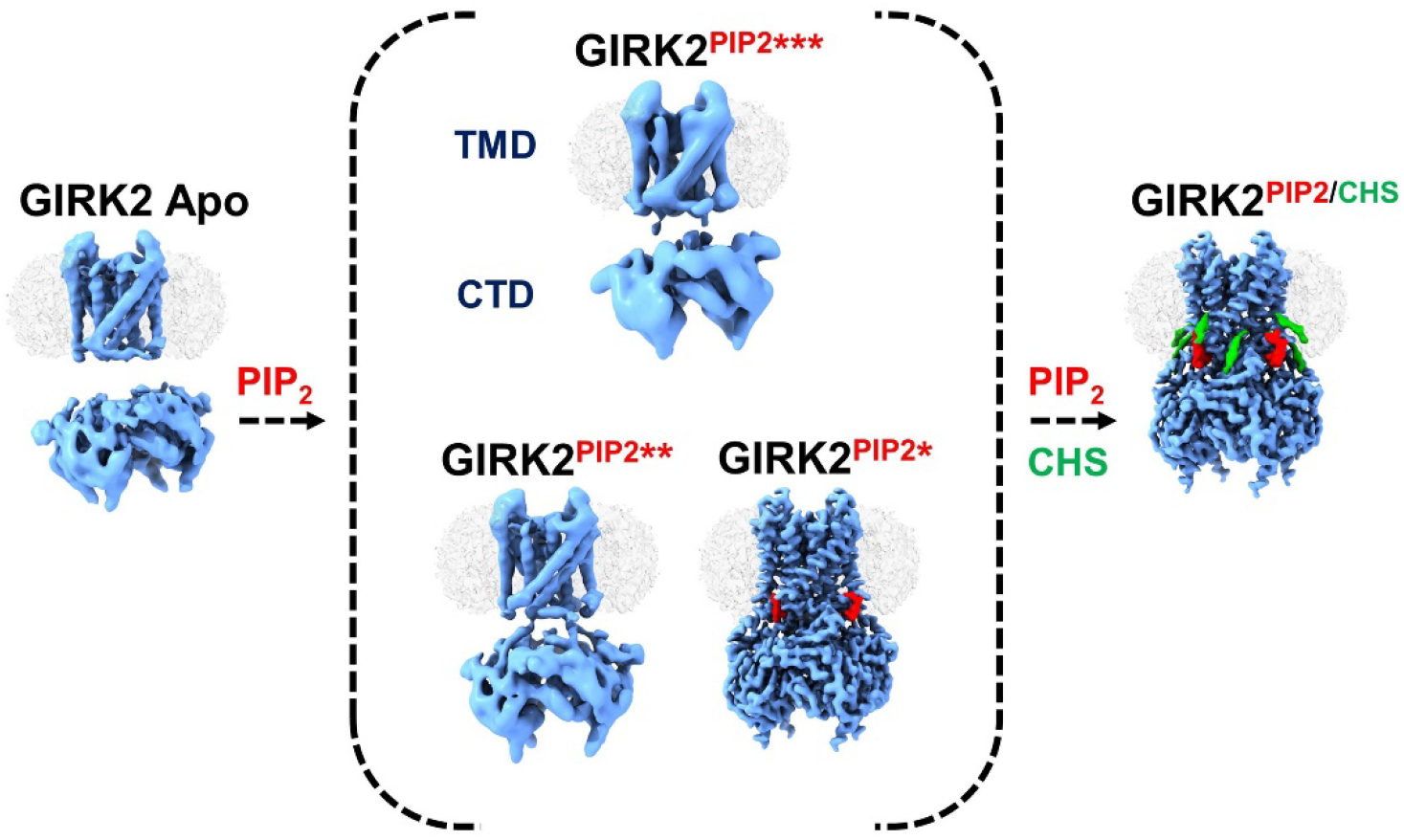
Different gating states for GIRK2 revealed by cryoEM. Structures with different lipid modulators suggest varying conformation of CTD with respect to TMD. The structures reveal novel cholesterol (green) binding sites adjacent to PIP_2_ (red) pocket. These molecules stabilize the TMD-CTD interface, thereby aiding in anchoring of CTD on to the membrane for further activation. The fraction of particles contributing to GIRK2 apo, GIRK2^PIP2^*, GIRK2^PIP2^**, GIRK2^PIP2^*** and GIRK2^PIP2/CHS^ maps as obtained from the cryoEM data processing is 22, 39, 27, 10 and 62%, respectively.

## DISCUSSION

The potential connection between elevated plasma cholesterol and increased risk for cardiac disease is well known^42^. Less well understood is the role of cholesterol in the brain. Nearly all cholesterol in the CNS is derived from *de novo* synthesis, and is therefore not influenced by dietary changes in cholesterol^43^. Nevertheless, changes in brain cholesterol have been implicated in numerous neurodegenerative diseases, such as Alzheimer’s disease (AD), Huntington’s disease, Parkinson’s disease, and Niemann-Pick disease^44^. For example, hypercholesterolemia causes impairment of dopamine signaling and psychomotor dysfunction in mice^45–48^. Increases in cholesterol levels have been shown to elevate beta-amyloid precursor protein levels in cholesterol-enriched lipid rafts^49^ and are associated with increased risk for AD^50^. Accordingly, there is a growing need to understand how cholesterol affects brain function.

### Cholesterol binding sites on membrane proteins

Cholesterol binding sites on membrane proteins were initially characterized using a variety of techniques, including electron spin resonance, photoaffinity labeling, molecular modeling and site-directed mutagenesis, which led to the identification of amino acid recognition motifs^15,37,51–54^. CRAC (Cholesterol Recognition/interaction Amino acid Consensus) is characterized by the amino acid sequence (L/V)-X_1–5_-(Y)-X_1–5_-(K/R)^55^, while CARC is oriented in the opposite direction (K/R)-X_1–5_-(Y/F/W)-X_1–5_-(L/V)^56^. The cholesterol consensus motif (CCM) is characterized by interactions with (W/Y)-(I/V/L)-(K/R) and (F/Y/R)^54^. By contrast, structures of membrane proteins (e.g., GPCRs and ion channels) determined in the presence of cholesterol/CHS have revealed interactions with “greasy hollows”, which refers to pockets in the transmembrane portion lined with hydrophobic residues^30,57,58^.

Docking studies and MD simulations identified putative cholesterol binding sites in GIRK channels that involved various amino acid residues^14,28,37^. In particular, two sites are very similar to the positions observed in the cryoEM structure (CHS1 and CHS2). The model for the first site implicated several residues involved in the CHS1 hydrophobic pocket, including L95, V99, and I175^14^. Additionally, the functional data suggest that V99 is involved in cholesterol binding. On the other hand, the second site is similar to the CHS2 pocket, as shown by functional and mutagenesis studies of several residues, including L86 and V101^14^. Thus, the putative cholesterol binding sites identified in our cryoEM structures are supported by results from previous mutagenesis studies.

### Mechanism of PIP_2_ gating in GIRK2 channels

Unexpectedly, we found the CTD is detached from the TMD in both the apo GIRK2 and a significant population of GIRK2^PIP2^ particles. By contrast, the CTD is engaged with the TMD in the structure of apo GIRK2 solved by X-ray crystallography^23^. One possibility for this difference could be due to lattice packing in X-ray crystal structures. Consistent with this idea, the CTD in the cryoEM structure of GIRK2^PIP2^* is ~ 1 Å lower from the membrane than in the X-ray structure of GIRK2^PIP2^ (PDBID:3SYA, **Extended Fig.S7a**). A detached CTD in the absence of PIP_2_ and other proteins (e.g. G proteins, SUR) may be a common feature of inward rectifiers, since the CTD is also detached in the apo Kir2.2 X-ray crystal structure^26^. Our cryoEM results suggest a plausible model for gating transitions in GIRK2 channels. First, PIP_2_ is clearly necessary for channel activation^13,38^. Second, the inter-domain region and the “wobbling” CTD are stabilized by the binding of four PIP_2_ molecules at the TMD-CTD interface, thus repositioning the stretched tether helix into a helical conformation and reducing the distance between Cα atoms of I195 and Q322 by 5-10 Å (**Fig. 4**). PIP_2_ most likely serves a crucial role in stabilizing the CTD interaction with the TMD region and promoting activation, which could explain why GIRK channels are not gated open by Gβγ, ethanol or cholesterol in the absence of PIP_2_.

In addition to the detached CTD, there are also changes in the coordination of PIP_2_ with GIRK2 between the cryoEM structures (GIRK2^PIP2/CHS^ and GIRK2^PIP2^*) and the X-ray crystal structure (PDBID:3SYA). The 1’ PO_4_ of PIP_2_ in the cryoEM structures interacts with the side chain of R92, whereas in the X-ray crystal structure it is coordinated by the amide backbone of R92 and the side chain is not resolved. This difference might be explained by the lower CTD in the cryoEM structures (**Extended Fig. 7a**). In addition, the 4’ PO_4_ interaction with K64 in 3SYA is not evident in the cryoEM structures. Interestingly, this pattern of engagement of PIP_2_ is similar to that observed with MD simulations of a constitutively open GIRK2 channel (K200Y)^59^, raising the possibility that the cryoEM structures of GIRK2 are closer to an open state. In support of this, the position of phosphate atom of 1’ PO_4_ of PIP_2_ is moved from the membrane towards cytoplasmic side by 3 Å and accompanied with ~ 3° rotation of the CTD, resulting in a wider pore opening at the inner-helix gate in the GIRK2^PIP2^* cryoEM structure, as compared to the crystal structure (**Extended Fig. S7**).

### Role of cholesterol on PIP_2_ mediated channel gating

Our cryoEM studies revealed one predominant conformation of GIRK2^PIP2/CHS^, whereas several additional conformers with detached CTDs were observed in the absence of CHS, suggesting that CHS may have a role in stabilizing the PIP_2_-bound GIRK2 with engaged CTD (**Fig. 5)**. These additional conformers of GIRK2^PIP2^ are likely due to the absence of cholesterol stabilization. Consistent with this observation, cholesterol stabilization of conformational states has been observed for several membrane proteins, in particular GPCRs, where the presence of cholesterol increases the thermal stability of the receptor and often lowers the energy barrier for agonist-induced conformational changes^60^. In addition, the binding of cholesterol in peripheral hydrophobic sites with no highly specific interactions is in agreement with the lack of large conformational changes on GIRK2 as a result of this engagement, implying its importance for primarily enhancing PIP_2_ binding. The increase in PIP_2_ affinity, as promoted by cholesterol, might increase the probability of the channel entering the open state. There is evidence that other allosteric modulators of channel activity such as alcohol and G proteins also act via increased PIP_2_ affinity^61^.

GIRK2 is relatively unique among members of the Kir family because it is potentiated in response to cholesterol rather than inhibited, as occurs with most other members of the Kir family^15^. As such, it presents a unique therapeutic target as well as a powerful model system to probe the gating mechanisms of Kir channel family members. The identification of cholesterol docking sites by structural studies allows us to probe the primary determinants of its interactions and integrate it with the results of computational docking, MD simulations and site-directed mutagenesis. Additionally, small molecules that weaken or strengthen the PIP_2_ affinity, potentially through regulating cholesterol binding, could be employed to modulate GIRK channel function in a manner tuned to respond to specific pathology.

## Supporting information

Extended Video S1_annot_small.MP4

## Acknowledgements

This work was supported in part by the National Institute on Alcohol Abuse and Alcoholism (AA018734) to PAS. We thank the Slesinger and Skiniotis laboratories for discussions on the experiments and E. Montabana for help with cryoEM data collection.

## Author Contributions

YKM, IWG, GS and PAS contributed to the design of the experiments; IWG and YZ prepared protein and IWG conducted functional studies; YKM prepared vitrified samples, collected and processed cryoEM data, and performed modeling. MJR implemented GemSpot; YKM and GS analyzed EM data; YKM, IWG, GS and PAS interpreted the results and wrote the manuscript

## ON-LINE METHODS

### Molecular Biology

A truncated *Mus musculus* GIRK2 cDNA (containing amino acids 52–380) linked in-frame with an HRV 3C protease site, green fluorescent protein (GFP) and a decahistidine (HIS10) tag (a generous gift from R. MacKinnon, The Rockefeller University, New York, NY) in pPICZ (ThermoFisher) was transformed into *Pichia pastoris* using electroporation (according to manufacturer protocols). Transformants were screened based upon Zeocin resistance (>1 mg/ml) and GFP emission. Highest expressing clones were selected for large-scale purification.

### Protein purification and reconstitution

All GIRK channels were expressed and purified in *P. pastoris* as described previously^13^. Briefly, the highest-expressing clone was grown in BMGY medium and induced in BMM medium containing 1% methanol. Cells were harvested, resuspended in buffer (50 mM HEPES, pH 7.4; 150 mM KCl; 1 mM TCEP; 1 mM AEBSF and Complete EDTA-free protease inhibitor tablets (Roche), flash frozen in liquid nitrogen, and stored at −80 °C. Frozen cells were lysed in a Mixer Mill (Retsch) 5-times for 3 minutes at 25 Hz and stored as powder at −80 °C until needed. The cell powder was solubilized in buffer containing 50 mM HEPES, pH 7.35; 150 mM KCl; 1 mM TCEP; 1mM AEBSF; Complete ULTRA EDTA-free protease inhibitor tablets (Roche) and either 2% (w/v) n-Dodecyl-β-D-maltoside (DDM; Anatrace) or 2% (w/v) DDM supplemented with 0.2% (w/v) Cholesteryl Hemisuccinate Tris Salt (Anatrace) with gentle stirring at 4 °C. Unsolubilized material was separated by centrifugation at 40,000 × g for 40 min at 4 °C and filtered. The supernatant was incubated with HISPur Cobalt charged resin (ThermoFisher) equilibrated in wash buffer (50 mM HEPES, pH 7.0; 150 mM KCl; 0.2% DDM or 0.2%DDM/0.02% CHS; 20 mM imidazole). The resin was subsequently washed in 10 column volumes (CV) of wash buffer, 5 CV containing 40 mM Imidazole, and eluted in buffer containing 300mM imidazole. The eluate was pooled, exchanged into imidazole-free buffer and digested overnight at 4 °C with HRV 3C protease, purified as described^62^ (a generous gift of Daniel Minor, UCSF, San Francisco, CA). The protein was subsequently concentrated and run on a Superdex-200 gel filtration column in buffer containing 20 mM TRIS-HCl pH 7.5, 150 mM KCl, 20 mM DTT, 3 mM TCEP, and 1 mM EDTA, 0.025% (w/v) DDM (anagrade) alone or with 0.0025% CHS. Fractions eluting at a volume consistent with the GIRK channel tetramer were pooled, concentrated and examined by SDS-PAGE and Coomassie blue staining.

Purified GIRK2 channels were reconstituted into lipid vesicles as described previously^13^. Briefly, a lipid mixture containing 1-palmitoyl-2-oleoyl-***sn***-glycero-3-phosphoethanolamine (POPE), 1-palmitoyl-2-oleoyl-***sn***-glycero-3-phospho-(1’-***rac***-glycerol) (POPG), and L-α-phosphatidylinositol-4,5-bisphosphate (Brain, PI(4,5)P_2_ (Porcine)) at mass ratios of 3:1:0.04 (POPE:POPG:PIP_2_) or 3:1 (POPE:POPG) was prepared, reconstituted in vesicle buffer (20 mM K-HEPES, pH 7.4; 150 mM KCl; 0.5 mM EDTA containing 35 mM CHAPS) and incubated with protein in detergent at a 1:200 protein: lipid ratio unless otherwise indicated. Where indicated, cholesterol (Ovine wool; Anatrace) was added to vesicles at a mole percentage of 5% or Cholesteryl Hemisuccinate (CHS; Anatrace) was incorporated at 2.5% mol percentage. Detergent was removed through sequential addition of Bio-beads SM-2 (Bio-rad). All phospholipids, cholesterol, and Brain PIP_2_ were purchased from Avanti Polar Lipids, Inc. Water-soluble PIP_2_ (diC_8_-PIP_2_) was purchased from Echelon Biosciences.

### Flux assay

The fluorescence-based flux assay for GIRK channel activity was performed as described previously^13,18^. Briefly, Liposomes were diluted 1:20 into flux buffer (20 mM Na-HEPES, pH 7.4; 150 mM NaCl; 0.5 mM EDTA) containing 5 μM of the H^+^ sensitive dye 9-Amino-6-chloro-2-μ methoxyacridine (ACMA)(Invitrogen). Fluorescence was measured using a Flexstation 3 microplate reader (Molecular Devices) with the following parameters: 410 nm excitation, 480 nm emission, 455 nm cutoff, medium PMT sensitivity, and sampling at 2 seconds. After a stable baseline fluorescence (150 s) was obtained, the H^+^ ionophore ***m***-chlorophenyl hydrazine (CCCP)(Sigma) was automatically added (1 μM final), then a second addition of vehicle or μ methanethiosulfonate hydroxyethyl (100 μM final; MTS-HE, Toronto Research Chemicals) was added 150 s later, followed 900 s later by a third addition with the K^+^ ionophore Valinomycin (100 nM final; Invitrogen), to determine the maximal K^+^ flux. GIRK2 channels are likely arranged in both orientations in the liposomes. However, we expect the channels oriented inside-out to support high K^+^ flux because of high Na^+^ in the flux buffer and high K^+^ in the liposome^13^. The percentage relative K^+^ flux was calculated by measuring the extent of quenching 10 s before VM addition, and dividing by the total quenching capacity of the liposomes normalized to the relative fluorescence units (RFU) 10 s before the addition of vehicle or MTS-HE. Liposome flux assay illustration created with Biorender.com

### CryoEM sample preparation, data collection and processing

The quality of purified samples was initially screened by negative stain EM using established protocols^39^. For cryoEM, 3.5 μl of GIRK2 at a concentration of 7-10 mg/ml was applied to glow-discharged Quantifoil Au1.2/1.3, 200 mesh grids, blotted and then plunge-frozen in liquid ethane using FEI Vitrobot. All cryoEM data were collected at 300 kV on a Titan Krios equipped with a Gatan K3 direct detection camera. Raw images were collected as movies, recorded at a magnification of 58,824 X corresponding to 0.85 pixel per Å at the specimen level. Each movie was recorded for 3 sec at 0.05 sec/frame with a total dose of 60 electrons/pixel and defocus values ranging from −0.8 to −2.2 μm.

The movies were motion-corrected and dose-weighted using Motioncor2^63^ and defocus values were determined by CTFFIND4^64^. Template-based particle picking, 2D/3D classification, and 3D refinements were performed using the Relion pipeline^65–68^. The 3D classes with both domains distinguishable were refined with C4 or C1 symmetry as shown in **Extended Figures S2, S4 and S6**, post processed with a mask encompassing both the domains, corrected for modulation transfer function (MTF) of K3 direct detection camera at 300 kV and sharpened with suitable B factor (**Extended Table S1 and S2**). The resolution of the maps reported here was estimated according to the 0.143 “gold-standard” Fourier Shell Correlation (FSC) criterion. The resolution of GIRK2^PIP2/CHS^, GIRK2^PIP2^*, GIRK2^PIP2^**, GIRK2^PIP2^***, GIRK2^PIP2^****, and apo GIRK2 was indicated globally as 3.5, 3.2, 4.8, 7.7, 7.7 and 4.8 Å, respectively. Local resolution was estimated using Relion. In apo GIRK2, the input particle stack for 3D classification was used for Relion multibody refinement analysis with TMD and CTD treated as separate bodies^69^(**Extended Fig. S5e and Extended Movie S1**).

### Modeling of protein and lipids in the cryoEM maps

The GIRK2 backbone and PIP_2_ at the TMD-CTD interface as determined in the X-ray crystal structure of GIRK2 (PDBID:3SYA) was initially fit as rigid-body into the EM maps using UCSF Chimera^70^ and subsequently interactively adjusted using COOT^41^. The GemSpot pipeline^40^ was used for modeling CHS into EM map followed by refinement in COOT^41^. The models in the target EM maps were refined further using Phenix real-space refinement^71^. Pore dimension in the model was analyzed using the HOLE program^72^. The figures in the manuscript were prepared using UCSF ChimeraX^73^

## Extended Supplemental Figures

**Extended Figure S1:**
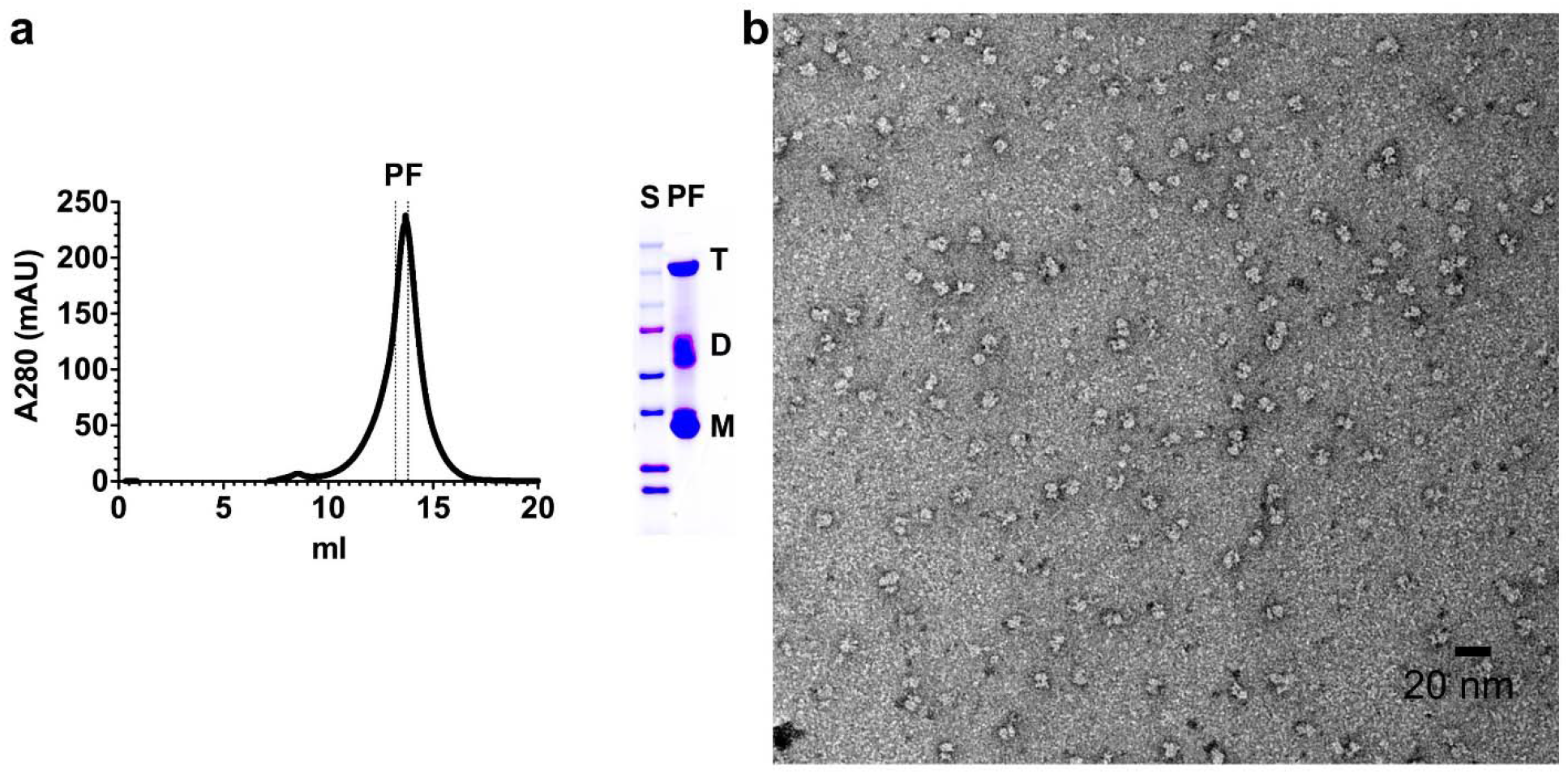
Purification and assessment of sample quality by negative staining electron microscopy. **(a)** Size exclusion chromatogram and Coomassie staining of purified GIRK2. The chromatogram (left panel) shows purification of mouse GIRK2 from *Pichia pastoris*. The absorbance is plotted as a function of elution volume. The major peak elutes at a volume consistent with the size of a ΔGIRK2 tetramer^1^. Peak fractions (PF) utilized in both functional and structural studies are indicated by dashed vertical lines. Coomassie blue staining (right panel) of pooled and concentrated peak fractions (PF) on a gradient SDS-PAGE protein gel shows three bands, one for tetramer (T), one for dimer (D), and one for monomer (M). **(b)** Micrograph of negative stained GIRK2. The image shows predominantly homogenous GIRK2 tetramers in detergent micelle.

**Extended Figure S2:**
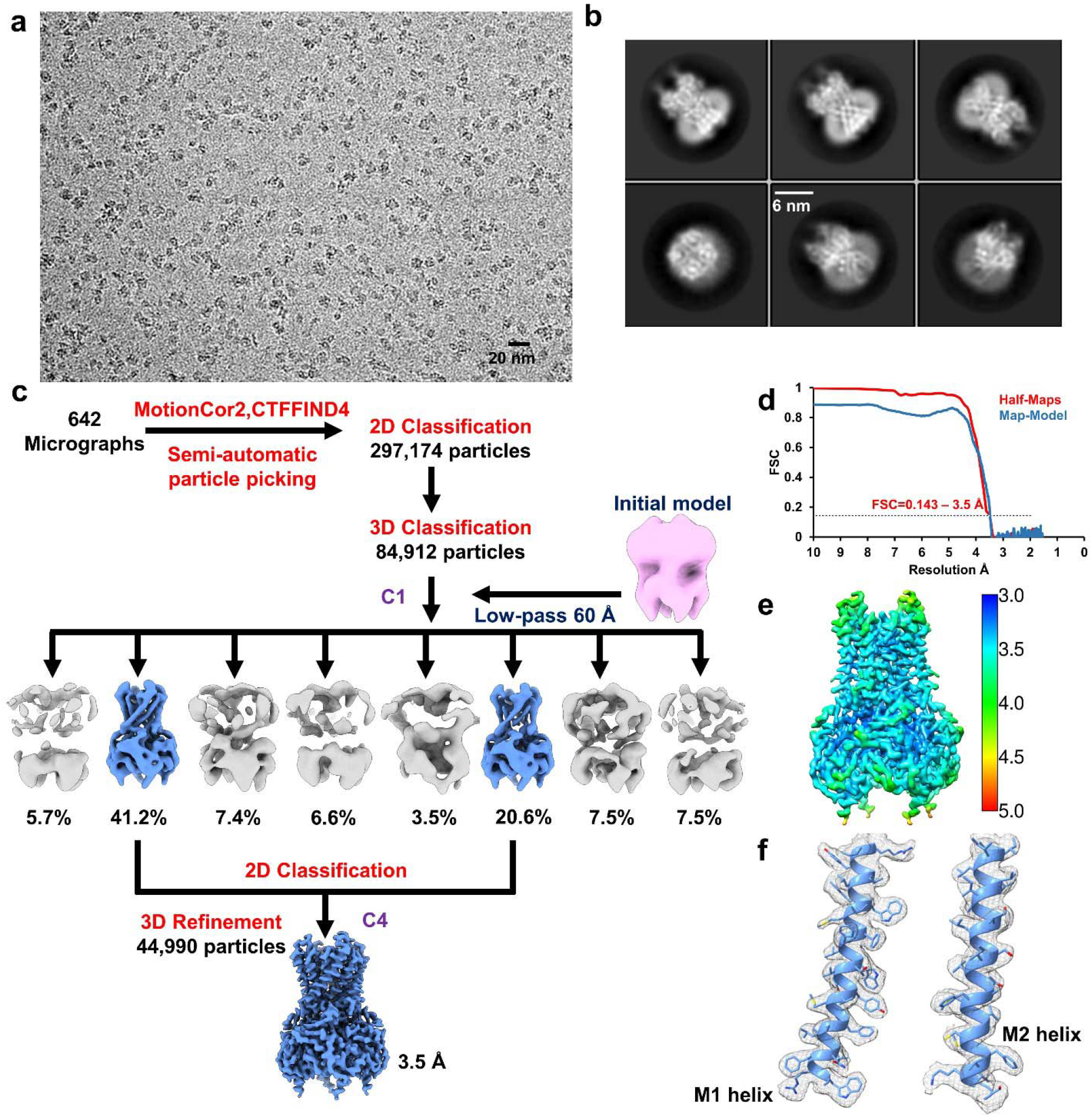
CryoEM data processing of GIRK2 in presence of modulators PIP_2_ and cholesteryl hemisuccinate. **(a)** and **(b)** are representative cryoEM image and 2D averages. **(c)** Data processing flow chart. **(d)** FSC curve, the dotted line indicating FSC at 0.143. **(e)** Local resolution estimate by Relion. **(f)** Fit of the transmembrane helices M1 and M2 into the 3.5 Å GIRK2^PIP2/CHS^ cryoEM map.

**Extended Figure S3:**
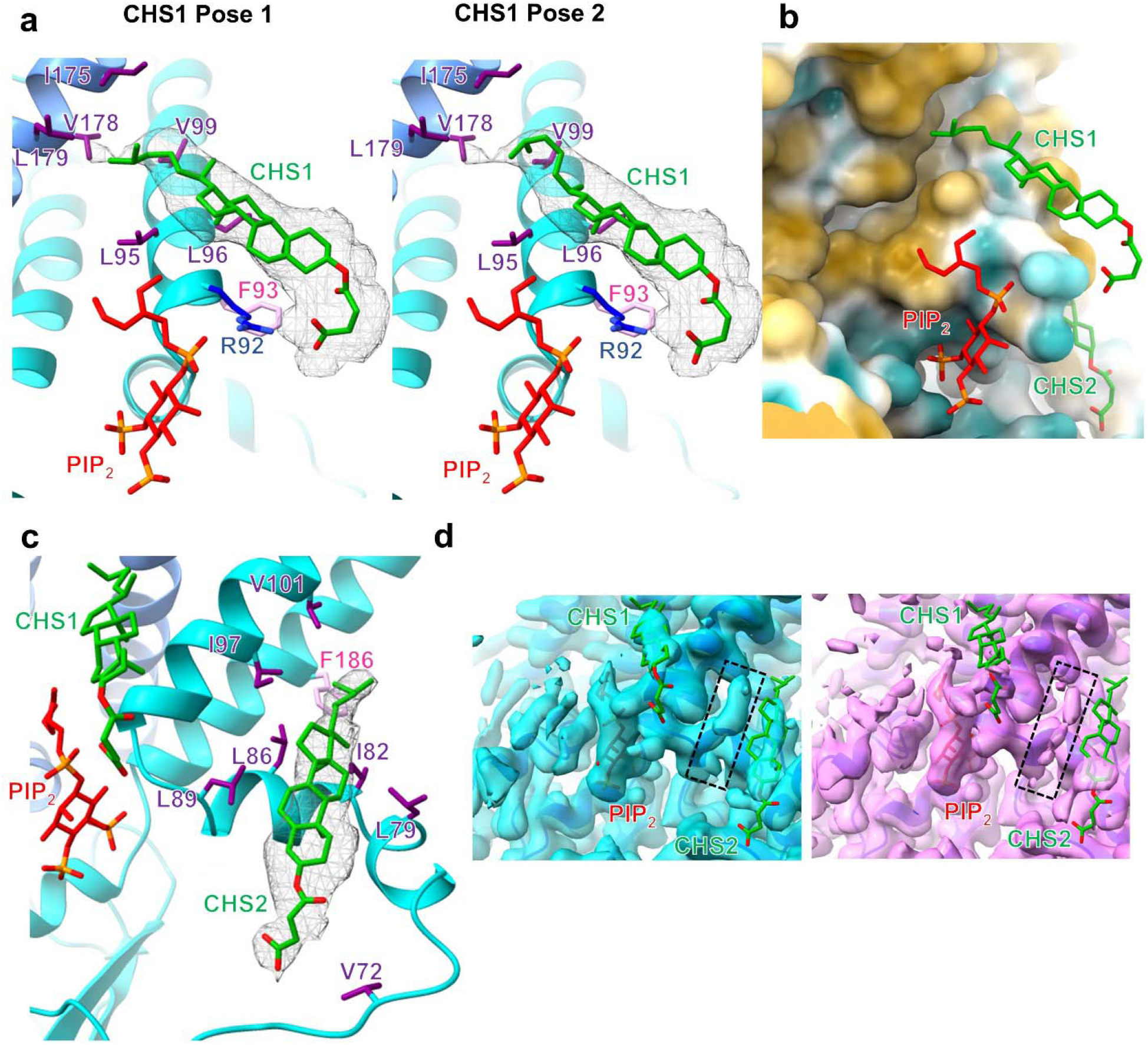
Modeling of cholesteryl hemisuccinate (CHS) in the cryoEM map. **(a)** Modelling of CHS1 (green) in GIRK2^PIP2/CHS^ cryoEM map suggests poses with subtle differences in the orientation of iso-octyl tail. The density of CHS1 is contoured at a threshold level of 0.004 in ChimeraX. **(b)** Illustration of residues around CHS1 colored based on hydrophobicity (dark cyan – most hydrophilic, yellow – most hydrophobic) in ChimeraX, CHS1 is bound in a hydrophobic environment. **(c)** CHS2 (green) is also surrounded by hydrophobic residues including F186 (pink), V72, L79, I82, L86, L89, I97 and V101 (purple). The density of CHS2 is contoured at a threshold level of 0.0073 in ChimeraX. **(d)** Comparison of GIRK2^PIP2/CHS^ (cyan, contoured at a threshold level of 0.0115 in ChimeraX) and GIRK2^PIP2^* (violet, contoured at a threshold level of 0.0095 in ChimeraX) maps, the position of CHS (green) and PIP_2_ (red) molecules are as in GIRK2^PIP2/CHS^ structure. Densities observed in GIRK2^PIP2/CHS^ map that were annotated as putative CHS sites are absent in GIRK2^PIP2^* map. Other poorly resolved densities near PIP_2_ consistent in both maps (highlighted one of them in black box) might correspond to other putative CHS or phospholipid binding sites.

**Extended Figure S4:**
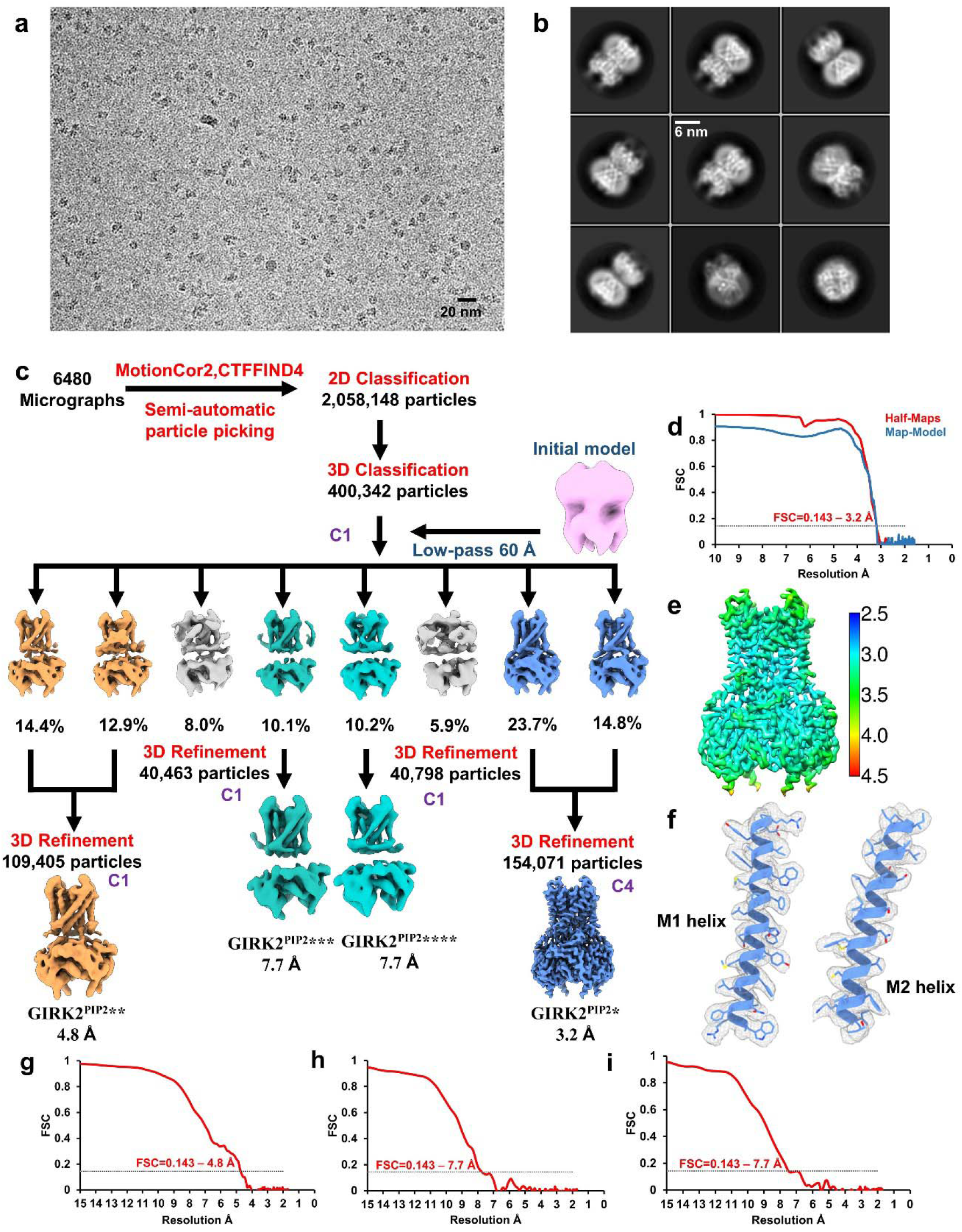
CryoEM data processing of GIRK2 in presence of PIP_2_. **(a)** and **(b)** are representative cryoEM image and 2D averages. **(c)** Data processing flow chart. **(d)** FSC curve of GIRK2^PIP2^*, the dotted line indicating FSC at 0.143. **(e)** Local resolution estimate of GIRK2^PIP2^* by Relion. **(f)** Fit of the transmembrane helices M1 and M2 into the 3.2 Å GIRK2^PIP2^* cryoEM map. **(g)** FSC curve of GIRK2^PIP2^**, the dotted line indicating FSC at 0.143. **(h)** FSC curve of GIRK2^PIP2^***, the dotted line indicating FSC at 0.143. **(i)** FSC curve of GIRK2^PIP2^****, the dotted line indicating FSC at 0.143.

**Extended Figure S5:**
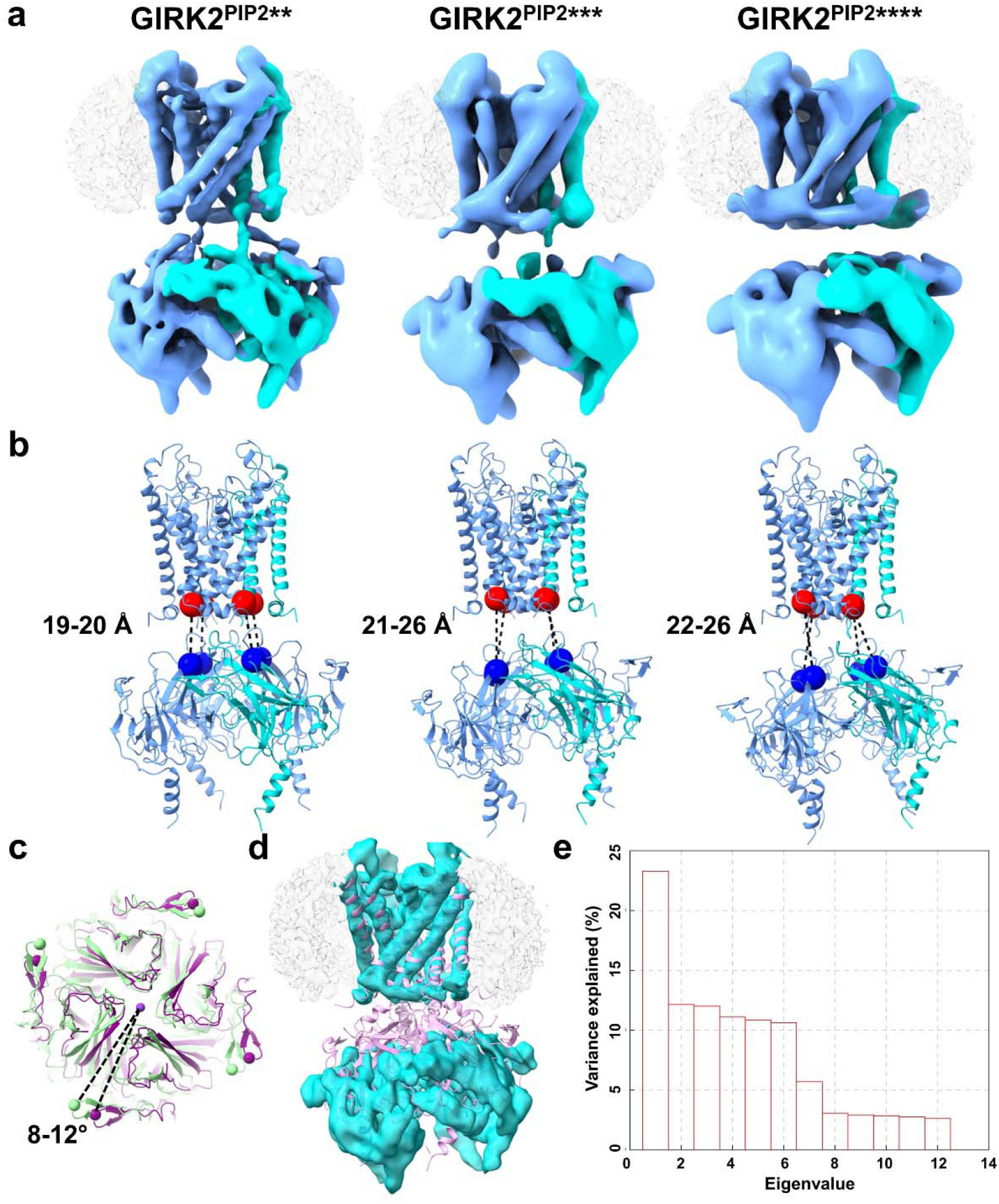
Evaluation of CTD detachment and twist in different structures. **(a)** CryoEM maps of GIRK2^PIP2^**, GIRK2^PIP2^*** and GIRK2^PIP2^**** with varying conformation of CTD with respect to TMD, one of the subunits is highlighted in cyan. **(b)** In these structures, CTD detachment is assessed by evaluating the distance between Cα atoms of residues I195 (denoted by red sphere) and Q322 (denoted by blue sphere). **(c)** Comparison of CTD twist of GIRK2^PIP2^** (purple) with GIRK2^PIP2^* (green) assessed by the angle of rotation of Cα atom of G347 (represented in sphere) around the four-fold axis after aligning the TMD near selectivity filter. **(d)** GIRK2 apo X-ray crystal structure (PDBID: 3SYO) is docked into GIRK2 apo cryoEM map. In contrast to the apo cryoEM map, the CTD in the crystal structure is engaged to the membrane. **(e)** Variance (%) for different eigenvalues obtained by multibody refinement of GIRK2 apo with TMD and CTD as separate bodies. A movie depicting the movements between TMD and CTD in 3D models for principal component 1 is provided as **Extended Movie S1**.

**Extended Figure S6:**
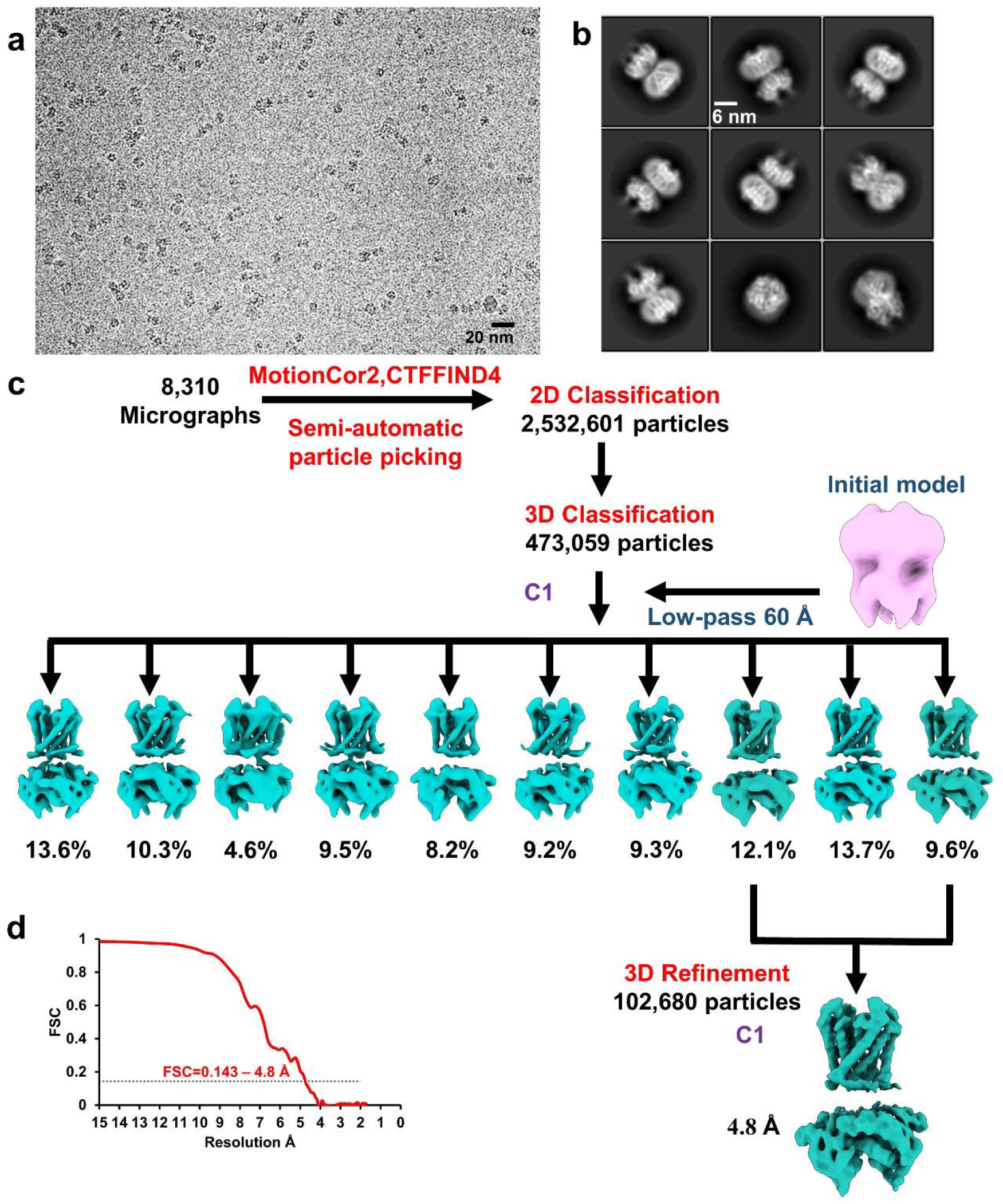
CryoEM data processing of apo GIRK2. **(a)** and **(b)** are representative cryoEM image and 2D averages. **(c)** Data processing flow chart. **(d)** FSC curve, the dotted line indicating FSC at 0.143.

**Extended Figure S7:**
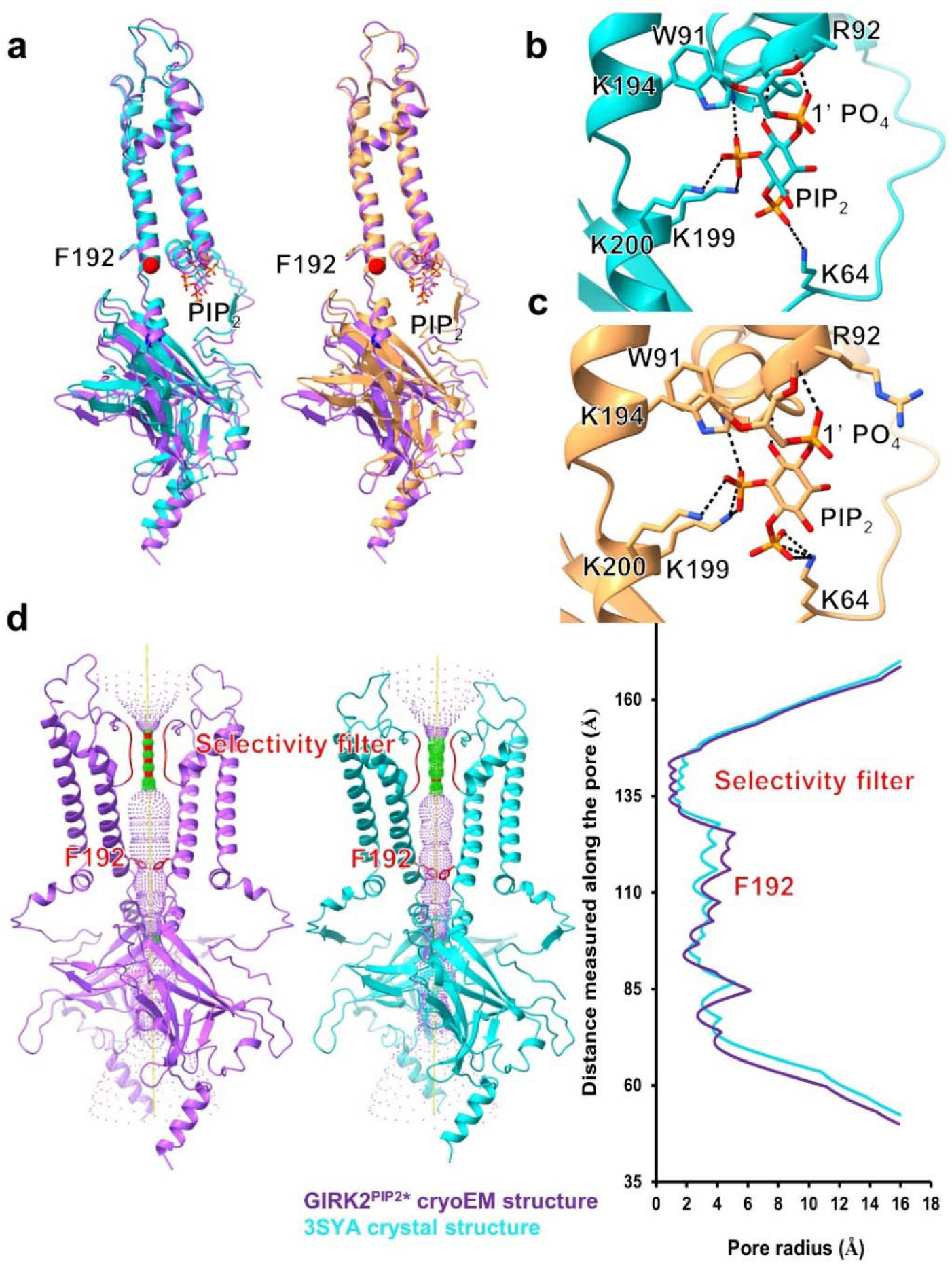
Comparison of PIP_2_ coordination, CTD and pore in cryoEM and X-ray crystal structures. **(a)** Comparison of PIP_2_ binding pocket and CTD in GIRK2^PIP2^* (violet), crystal structures with only PIP_2_ (PDBID:3SYA, blue) and with Gβγ and PIP_2_ (PDBID:4KFM, orange) after aligning the TMD near selectivity filter. There are subtle differences in the position of PIP_2_, CTD and inner-helix gate residue F192. Compared to crystal structure with only PIP_2_ (3SYA), the positions of phosphate atom of 1’ PO_4_ of PIP_2_ and CTD (evaluated by distance between Cα atoms of I195, red sphere and Q322, blue sphere) in GIRK2^PIP2^* cryoEM structure are moved away from the membrane towards the cytoplasmic side by ~ 3 and 1 Å, respectively, and accompanied with ~ 3° rotation of CTD around the four-fold axis. **(b)** and **(c)** are PIP_2_ coordination in crystal structures PDBID:3SYA (blue) and PDBID:4KFM (orange), respectively. In crystal structures, 1’PO_4_ of PIP_2_ is coordinated by backbone amide of R92 in contrast to the side chain as seen in cryoEM structures. **(d)** Pore comparison using HOLE software for cryoEM structure GIRK2^PIP2^* (violet) and X-ray crystal structure PDBID:3SYA (blue). The pore radius around the inner-helix gate (F192) is wider in the cryoEM structure when compared to 3SYA.

**Extended Movie S1:** Movie depicting the models obtained from multibody refinement of GIRK2 apo with transmembrane (TMD) and cytoplasmic (CTD) domains as two separate bodies.

Filename: Extended Video S1_annot_small.MP4

**Extended Table S1:**
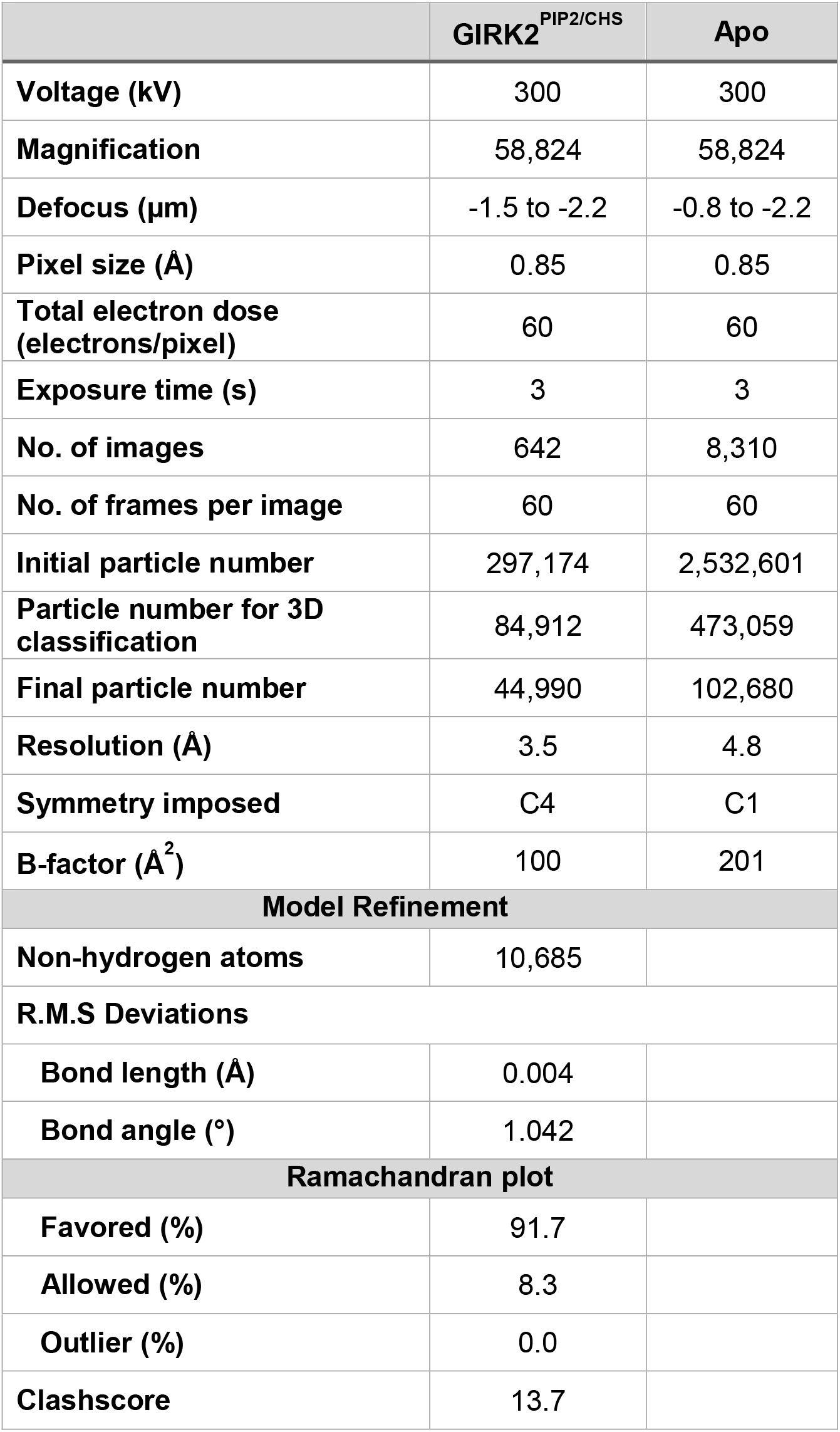
Data collection and processing statistics of GIRK2^PIP2/CHS^ and apo structures.

|  | **GIRK2<sup>PIP2/CHS</sup>** | **Apo** |
| --- | --- | --- |
| **Voltage (kV)** | 300 | 300 |
| **Magnification** | 58,824 | 58,824 |
| **Defocus (μm)** | -1.5 to -2.2 | -0.8 to -2.2 |
| **Pixel size (Å)** | 0.85 | 0.85 |
| **Total electron dose (electrons/pixel)** | 60 | 60 |
| **Exposure time (s)** | 3 | 3 |
| **No. of images** | 642 | 8,310 |
| **No. of frames per image** | 60 | 60 |
| **Initial particle number** | 297,174 | 2,532,601 |
| **Particle number for 3D classification** | 84,912 | 473,059 |
| **Final particle number** | 44,990 | 102,680 |
| **Resolution (Å)** | 3.5 | 4.8 |
| **Symmetry imposed** | C4 | C1 |
| **B-factor (Å<sup>2</sup>)** | 100 | 201 |
| **Model Refinement** |  |  |
| **Non-hydrogen atoms** | 10,685 |  |
| **R.M.S Deviations** |  |  |
| **Bond length (Å)** | 0.004 |  |
| **Bond angle (°)** | 1.042 |  |
| **Ramachandran plot** |  |  |
| **Favored (%)** | 91.7 |  |
| **Allowed (%)** | 8.3 |  |
| **Outlier (%)** | 0.0 |  |
| **Clashscore** | 13.7 |  |

**Extended Table S2:**
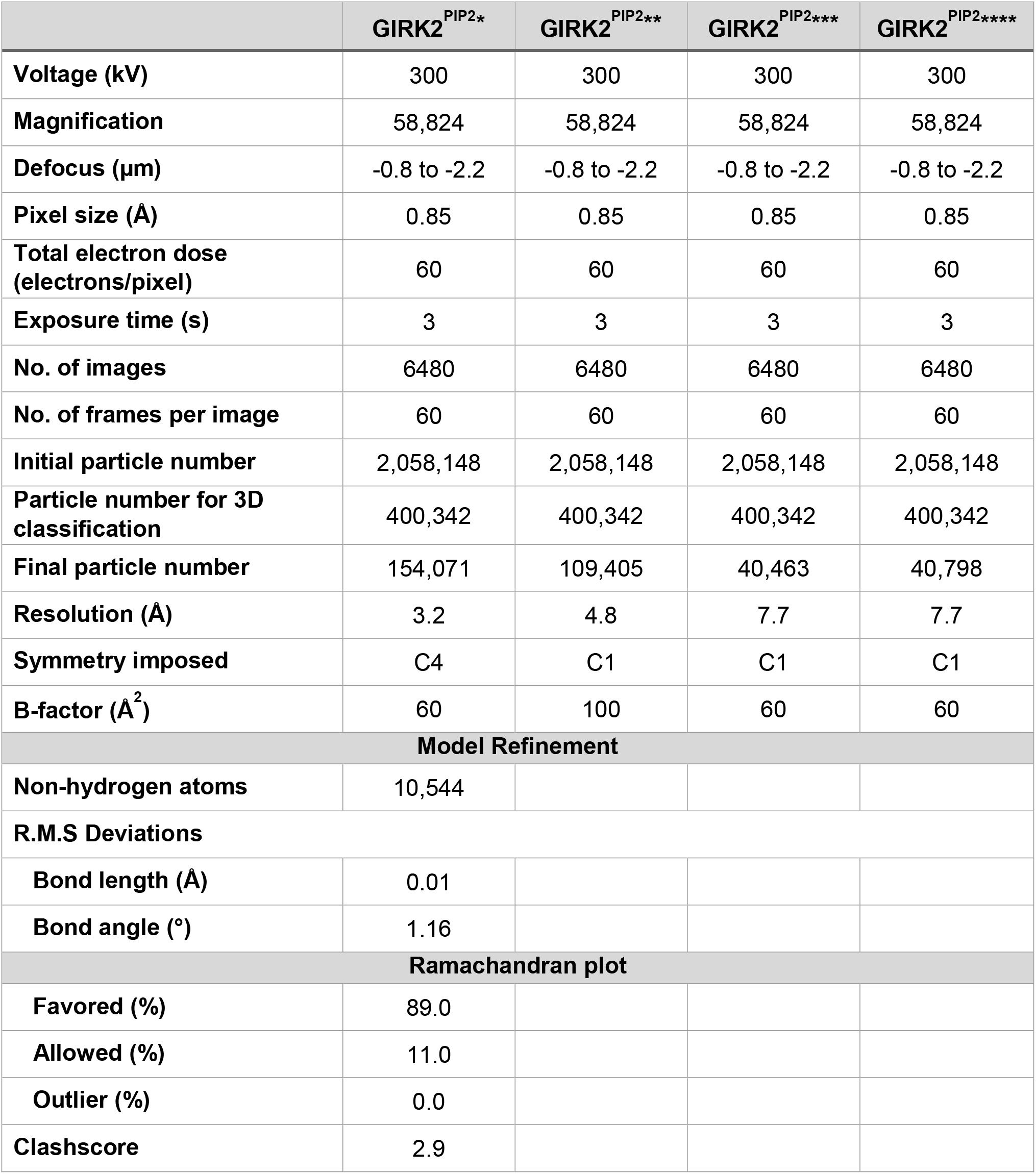
Data collection and processing statistics of GIRK2^PIP2^ structures.

